# Temporal dynamics of color polymorphism and hybridization in *Colias* butterflies

**DOI:** 10.1101/2025.05.25.656022

**Authors:** Joshua P. Jahner, Matthew L. Forister, Thomas L. Parchman, Shelby M. Burdo, S. Eryn McFarlane, C. Alex Buerkle, Chris C. Nice, James A. Fordyce, Arthur M. Shapiro

**Author notes:** Corresponding author: Department of Botany, University of Wyoming, Laramie, WY, USA.

## Abstract

Investigations into the genetic basis of color polymorphism have played a key role in our understanding of genetic architecture and the evolution of mating systems. Sulphur butterflies (*Colias*) have been models in this field, but also contain unsolved puzzles with respect to species boundaries and hybridization. We surveyed genomic variation across five years in a location where phenotypic intermediates between *Colias eurytheme* and *C. eriphyle* are observed, but ancestry variation of potential hybrids has not been quantified. Our results reveal individuals with hybrid ancestry roughly in proportion to the frequency of individuals of intermediate phenotype recorded in the wild. Individuals with hybrid ancestry were predominantly those with intermediate morphologies, but morphologically intermediate individuals were not always of hybrid origin, which raises alternative possibilities for the origin and maintenance of color variation in the system. Genomic regions differentiated between species are largely located on the Z chromosome, and we find more candidates on autosomes than in another *Colias* contact zone. The dynamics of hybridization in this system are highly variable through time, suggesting fertile avenues for future study into the maintenance of species boundaries in the face of temporally-variable, climatically influenced, and pervasive hybridization.

## Introduction

Understanding color variation within and among species has long been a goal that unites various disciplines within ecology, evolutionary biology, and animal behavior (Cuthill *et al*., 2017). Insects, and butterflies in particular, provide some of the most spectacular examples of coloration that are also associated with a wealth of natural history knowledge relative to most other insect groups for which such information is often lacking (Monteiro & Prudic, 2010). From a genetic perspective, coloration in butterflies has also been of particular interest because it has been, in many cases, linked to large effect loci on sex chromosomes (Ellers & Boggs, 2003; Sahara *et al*., 2012; Brien *et al*., 2019). Sex-linked traits can potentially be important to reproductive isolation because, in hybrids, selection against hemizygous, deleterious alleles in the heterogametic sex is expected to be more effective at limiting gene flow than selection at diploid sites. The butterflies *Colias eurytheme*, *C. philodice*, and *C. eriphyle* are known to hybridize and also rely on a sex-linked mating signal that is based on differences in color and UV reflectance (Rutowski *et al*., 2005; Ficarrotta *et al*., 2022). A recent examination of 24 males from a single *C. eurytheme* and *C. philodice* population in the eastern United States reported extreme divergence across the entire Z chromosome (where the gene associated with UV reflectance resides), yet little evidence of differentiation elsewhere in the genome, resulting in the largest known ratio of sex chromosome to autosome divergence (Ficarrotta *et al*., 2022). Hanly *et al*. (2023) followed this study with quantitative trait mapping of lab generated hybrids, finding two quantitative trait loci on the Z and chromosome 18 that explain roughly 70% of the variation in wing color. Here we offer a complementary study using genome scans of wild-caught individuals across multiple years in the context of a long-term hybrid zone study.

Hybridization in North American *Colias* butterflies has been described as emblematic of the evolutionary puzzle of species boundaries because hybridization is rampant yet parental taxa remain morphologically distinct (Shapiro, 2012). Taxonomy in this group has been controversial, especially with respect to the status of *C. philodice* and *C. eriphyle*. The latter is often treated as either synonymous with or a subspecies of the former, but we refer to *C. eriphyle* as a specific entity in this paper, following Wheat & Watt (2008); also see Zhang *et al*. (2019) where the two taxa plus *C. eurytheme* are treated as nominal entities in a phylogenetic analysis. Not only is hybridization common among these species, but it is also highly variable across time and space. Temporal dynamics are sensitive to climatic shifts including drought. Jahner *et al*. (2012) found that much of the variation in the frequency of phenotypically-identified hybrids between *C. eurytheme* and *C. eriphyle* could be predicted by abiotic conditions. In particular, previous-year temperature affected the abundance of parental species, likely mediated by host plant quality, which in turn affected the abundance of hybrids (Jahner *et al*., 2012). In space, much of the complexity derives from the introduction of alfalfa (*Medicago sativa*) to North America within the last 200 years. All three butterfly species use the exotic host and have expanded in various ways as a consequence of that colonization, such that there are now few places where non-hybrid populations can be found of any of the three species. One of these places is Mediterranean-climate regions of California, which forms the backdrop for the present study. The most recent genetic study in the western United States found that phenotypically-identified hybrid specimens were not identifiable as genetic hybrids (Dwyer *et al*., 2015). That finding added to the evolutionary puzzle of *Colias* by suggesting that apparent hybrids between the two taxa, *C. eurytheme* and *C. eriphyle*, are actually color morphs within *C. eriphyle* that could be the result of parallel evolution. Dwyer *et al*. (2015) used AFLP markers and noted the need for reanalysis using modern genomic tools.

Given the spatial and temporal dynamism of geographic ranges and hybrid zones in *Colias*, it has been suggested that multi-year sampling might be needed to clarify the picture of species boundaries and hybridization (Shapiro, 2012). For example, differences in over-wintering requirements among the taxa mean many individual hybrid zones reform annually with one of the species immigrating each spring. It is precisely this biogeographic context that we analyze here, studying parental and phenotypic hybrids across five consecutive years in a single well-studied location, asking the following questions. (1) Do the two taxa (*C. eurytheme* and *C. eriphyle*) hybridize, and are morphologically-identified intermediate individuals actually genetic hybrids, or are apparent (morphological) hybrids variants within one or both of the parental taxa as suggested by Dwyer *et al*. (2015)? (2) What is the genetic architecture of color polymorphism in this system? With respect to the second question, the expectation from the study of hybrids between *C. eurytheme* and *C. philodice* (Ficarrotta *et al*., 2022; Hanly *et al*., 2023) is overwhelming dominance of the sex chromosome in both color determination and differentiation of species.

## Materials and methods

### Sequencing and bioinformatics

Adult butterflies were sampled from a single alfalfa field in Sierra Valley, California during the active flight window roughly every two weeks from 1983 – 2004 (Fig. 2; Jahner *et al*., 2012). During each sampling event, butterflies were collected indiscriminately by color and sex either for two person hours or until a total of 100 individuals were netted. On average, 428 *C. eurytheme*, 130 *C. eriphyle*, and 31 hybrids were sampled per year from 1983 – 2004 (Jahner *et al*., 2012). For this research, we considered individuals from a five year window (1997 – 2001) that immediately followed a temporary extirpation of *C. eriphyle* from Sierra Valley (Fig. 2; Jahner *et al*., 2012). Specimens were classified by AMS as one of four color phenotypes (*C. eriphyle* = yellow; *C. eurytheme* = orange; hybrids = intermediate; alba females = white) (Fig. 1). The alba phenotype (white females; Fig. 1) has a single genetic origin that has been maintained in roughly one-third of *Colias* species via introgression and balancing selection (Tunström *et al*., 2023). Throughout, we refer to the parental species predominantly with scientific names (*C. eriphyle*, *C. eurytheme*), but also in some cases with colors (yellow, orange) when it facilitates interpretation; however, when individuals with hybrid or alba phenotypes are genetically assigned to one of the parental species, scientific names are used as subscripts (Hybrid_EU_, Hybrid_ER_, Alba_EU_, Alba_ER_), as explained further below.

**Figure 1:**
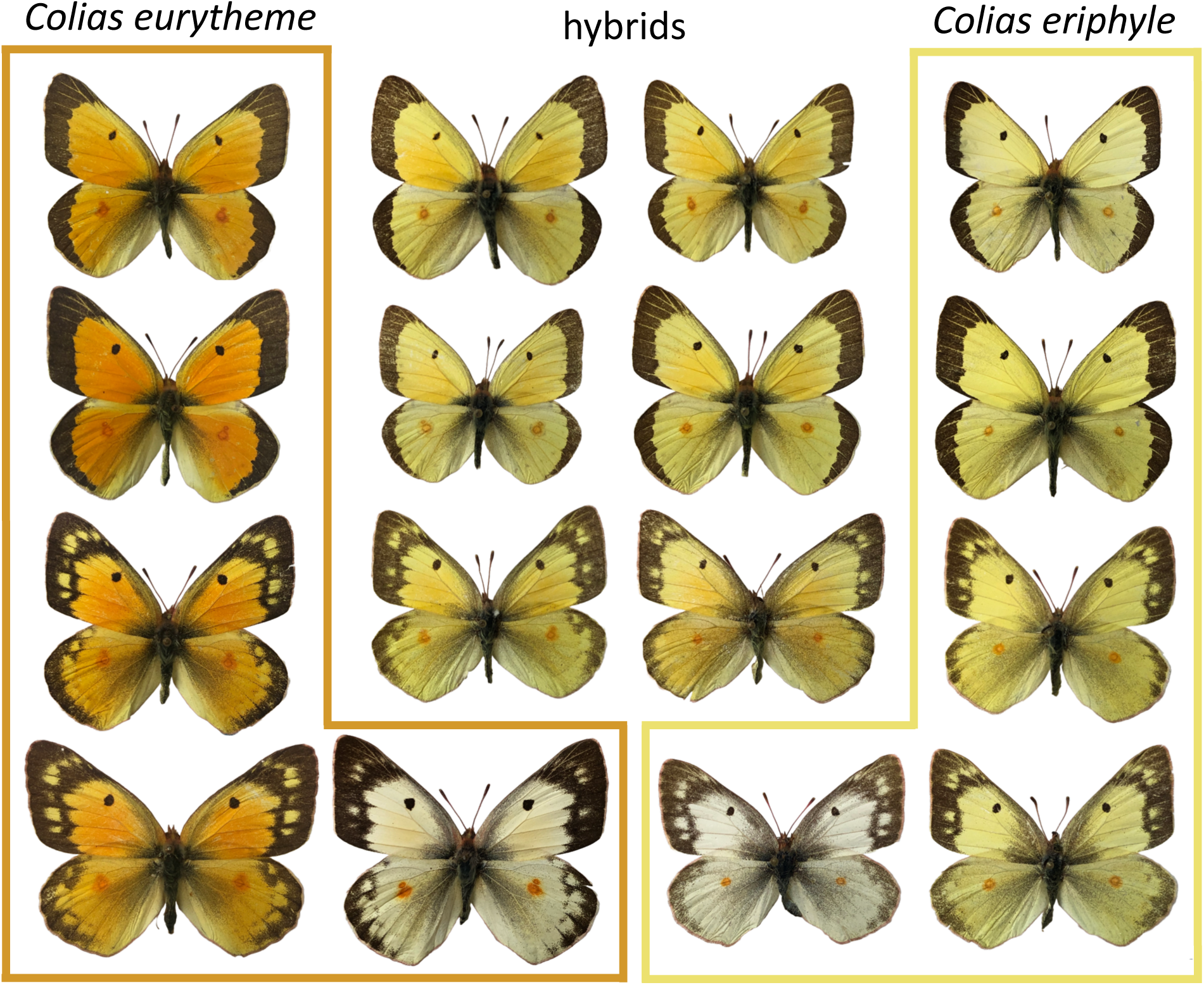
Parental and hybrid wing phenotypes (upper surfaces shown). *Colias eurytheme* are shown along the left side, surrounded by orange lines; *C. eriphyle* are along the right side, surrounded by yellow lines; six hybrid individuals are in the middle (note the orange coloration on the fore wings of the hybrid individuals). Males are in the top two rows, females in the bottom two rows; one alba (white) individual is shown for each of *C. eurytheme* and *C. eriphyle*.

**Figure 2:**
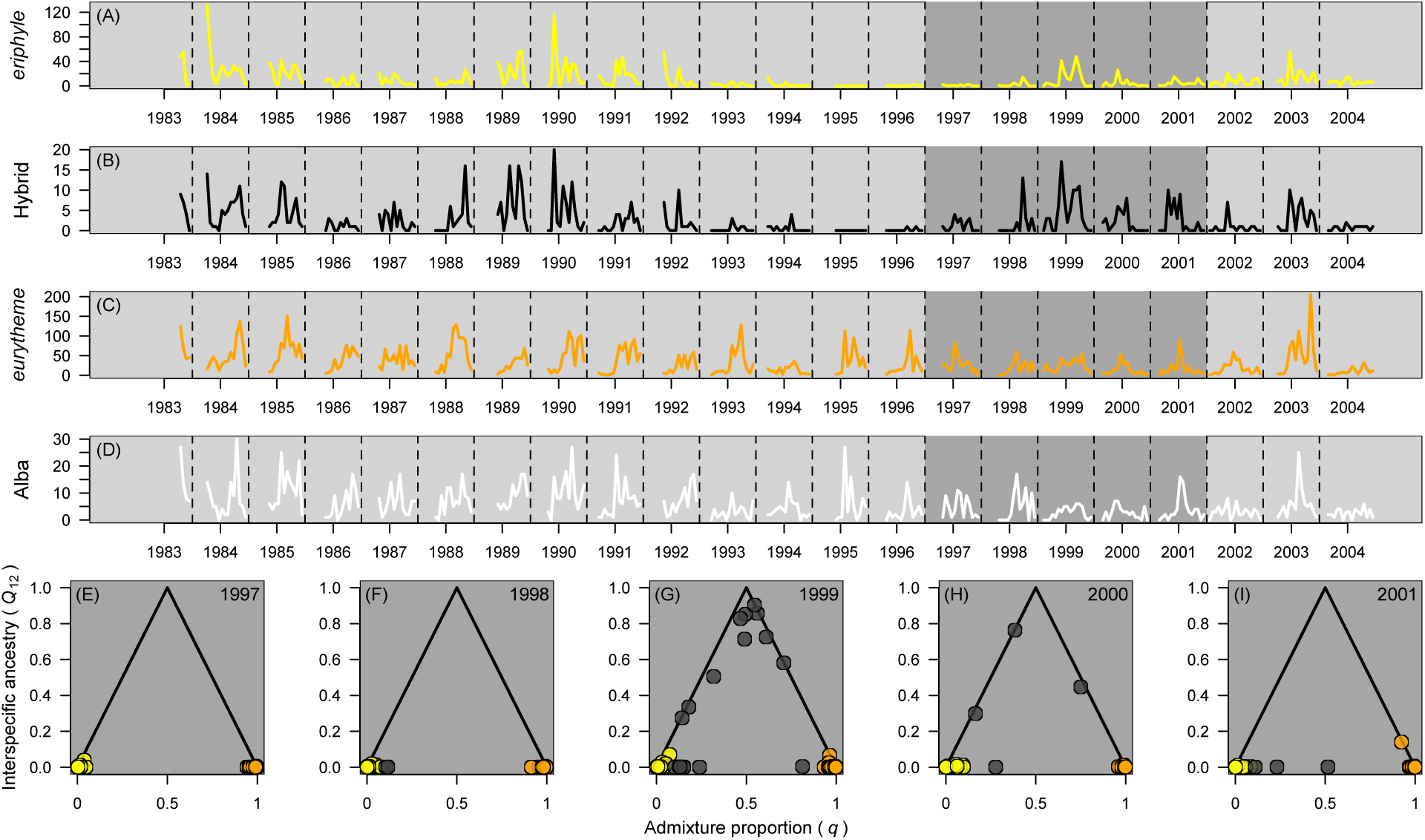
(**A–D**) The number of *C. eriphyle* (yellow), hybrid, *C. eurytheme* (orange), and alba (white) individuals sampled from 1983 – 2004 (data from Jahner *et al*., 2012). Only a subset of individuals from 1997 – 2001 were included in this study; those years are highlighted with darker gray in time series plots. **(E–I)** The frequency of hybridization differed across years. Individual ancestry classes were assigned based on estimates of admixture proportion (*q*) and interspecific ancestry (*Q*_12_) from entropy. Individuals with parental ancestry have *Q*_12_ = 0 and either *q* = 0 (*C. eriphyle*) or *q* = 1 (*C. eurytheme*); F_1_ hybrids have *q* = 0.5 and *Q*_12_ = 1; F_2_ hybrids have *q* = 0.5 and *Q*_12_ = 0.5 on average; and backcrosses have combinations of *q* and *Q*_12_ that reside on the solid black lines. Each point corresponds to an individual butterfly, and points are colored based on their admixture proportion (yellow: *q <* 0.1; black: 0.1 *< q <* 0.9; orange: 0.9 *< q*). See Fig. S1 for ancestry results for only male individuals.

All putative hybrids (*N* = 157) from 1997 – 2001 were sequenced, as well as 111 *C. eriphyle*, 137 *C. eurytheme*, and 75 alba (total *N* = 480; Table S1). For the latter three categories, individuals were deliberately selected from throughout the five year window to capture as much temporal variation as possible. DNA was extracted from the anterior third of the abdomen using Qiagen DNeasy Blood and Tissue kits (Qiagen Inc., Valencia, CA), and genotyping-by-sequencing (GBS) libraries were constructed following Parchman *et al*. (2012). Briefly, we restricted extracted DNA using the enzymes *Eco*RI and *Mse*I (one expected cut every 4,096 and 256 base pairs, respectively), ligated a barcode indentifier (unique to each individual butterfly) to each fragment, and amplified fragments using PCR. Libraries were size-selected to include DNA fragments 350 – 450 base pairs in length using a Pippin Prep (Sage Science, Beverly, MA), and sequenced on two Illumina HiSeq 2500 lanes at the University of Texas Genomic Sequencing and Analysis Facility (Austin, TX).

Contaminant reads (e.g., *E. coli*; PhiX) were removed from the dataset using the tapioca pipeline (https://github.com/ncgr/tapioca) and bowtie_db2 (Langmead & Salzberg, 2012). Reads were assigned to individuals based on unique barcodes that were ligated onto sequences during library prep, and then split into separate individual fastq files using custom Perl scripts. Reads were aligned to the *C. croceus* chromosome-level genome assembly (ilColCroc2.1; GCA_905220415; Ebdon *et al*., 2021) using the *aln* and *samse* algorithms of bwa v0.7.17 (Li & Durbin, 2009) specifying a maximum edit distance of 4. Sorted sam files were converted to binary format (bam) and variant sites were called using bcftools v1.9 and samtools v1.9 (Li *et al*., 2009) specifying minimum base, site, and mapping qualities of 20, a minimum genotype quality of 10, and a max depth of 100.

A final set of single nucleotide polymorphisms (SNPs) were retained for downstream population genetic analyses following a series of filtering steps performed with vcftools v0.1.14 (Danecek *et al*., 2011). Based on preliminary data exploration, a single individual that did not sequence well was removed from the dataset. Next, sites were retained if the minor allele frequency was greater than 0.02 and the proportion of individuals with missing data was less than 0.7. Additionally, only biallelic sites and a single variant per contig (87–89 base pairs long) were retained. In an effort to exclude paralogous sites, variants were removed if the maximum average depth across individuals was greater than 35 (*N* = 144; 1.4% of loci) and if *F* _IS_ was less than –0.5 (*N* = 6).

### Quantifying individual ancestry

The ancestry complement model of entropy v2.0 (Gompert *et al*., 2014; Lindtke *et al*., 2014; Shastry *et al*., 2021) was used to estimate two hybridization metrics – admixture proportion (*q*) and interspecific ancestry (*Q*_12_) – that are useful for assigning individuals to different ancestry classes. Specifically, *q* represents the proportion of the genome derived from one of two different parental ancestries (only two parental ancestries were considered in this study), whereas *Q*_12_ reflects the proportion of the genome that has a combination or complement of two ancestries at a locus. Expected values of *q* and *Q*_12_ were used to assign each individual to an ancestry category: individuals with parental ancestry have *Q*_12_ = 0 and *q* = 0 or 1; F_1_ hybrids have *Q*_12_ = 1 and *q* = 0.5; F_2_ hybrids have *Q*_12_ = 0.5 and *q* = 0.5 (on average); and backcrossed individuals reside on lines connecting F_1_ hybrids and parental individuals (Gompert *et al*., 2014; Lindtke *et al*., 2014; Shastry *et al*., 2021). Five replicate entropy chains were run for 250,000 MCMC iterations, with a burn-in of 50,000 and a thinning interval of 40. Chain mixing and convergence were assessed based on effective sample size and potential scale reduction factor (*R*^^^).

Because we were particularly interested in divergence on the Z chromosome (Ficarrotta *et al*., 2022; Hanly *et al*., 2023), we also ran an additional model for males only (*N* = 242; *C. eurytheme* = 68; *C. eriphyle* = 59; hybrid = 115) using the same specification as above. We ran this analysis to safeguard against any potential complications that could arise from treating all individuals as diploid during variant calling, even though females are haploid for the Z chromosome (i.e., ZW).

### The genomic landscape of color-associated differentiation

In addition to the ancestry metrics described above, entropy also generates genotype probabilities for each individual at each locus. Genotype probabilities range from 0 – 2 (the two alternative homozygous states; heterozygotes = 1) and are valuable for low-coverage sequencing datasets because they incorporate inherent uncertainty that arises during sequencing and genotype calling (Nielsen *et al*., 2011; Buerkle & Gompert, 2013; Fumagalli *et al*., 2013). All population genetic analyses were based on genotype probabilities.

To predict wing color phenotypes (orange, yellow, alba, hybrid) from the sequencing data, a series of random forest models were obtained using the randomForest package in R v4.2.1 (Liaw & Wiener, 2002; R Core Team, 2022). All orange and yellow individuals were classified based on wing morphology. For morphologically alba and hybrid individuals, models either grouped all individuals together (Alba_ALL_; Hybrid_ALL_) or split individuals based on admixture proportion (*q*) from entropy (Alba_EU_ or Alba_ER_; Hybrid_HY_, Hybrid_EU_, or Hybrid_ER_). In all color models, phenotype was predicted by the full matrix of genotype probabilities. Additionally, to identify potential genetic differences across years, a final set of three models was run for all individuals, morphologically orange individuals, and morphologically yellow individuals. Model performance was assessed using the out of bag (OOB) error percentage, and the predictive performance for individual loci was determined using the mean decrease in the Gini Index.

As a complementary approach to random forest, the Bayesian sparse linear mixed model (BSLMM) of gemma v0.98.4 (Zhou *et al*., 2013) was used to predict color phenotype from genotype. This model identifies a sparse subset of predictors (i.e., SNPs) that explain phenotypic variation after accounting for a relatedness matrix among all individuals. Additionally, the model estimates a posterior inclusion probability (PIP) for each predictor, which is the probability that a given locus is included in the sparse subset of predictors, and therefore associated with the phenotype of interest. Because the random forest models only identified meaningful differences among orange and yellow individuals, a BSLMM was only performed for the morphologically orange versus yellow comparison. The model was run for 25,000,000 iterations with a burn-in of 10,000,000 and a thinning interval of 10,000.

To characterize differentiation and divergence between morphologically orange and yellow individuals, locus-specific pairwise *F* _ST_ (Hudson *et al*., 1992) and *π*_b_ (Charlesworth, 1998) were calculated based on allele frequencies (i.e., mean population genotype probability per color / 2). *F* _ST_ is a relative measure of differentiation between groups that is dependent on within-group levels of genetic diversity. Because high values of *F* _ST_ can manifest from both high differentiation and low within-group diversity, this metric has limited utility to identify regions of the genome associated with glittery (*sensu* Rockman, 2012) traits of interest (e.g., butterfly wing colors) (Charlesworth, 1998; Cruickshank & Hahn, 2014; Ravinet *et al*., 2017). However, *F* _ST_ may have some utility in cases where gene flow between populations is sufficiently high (e.g., Martin *et al*., 2013). In contrast, *π*_b_ is an absolute measure of divergence that is independent of within-group diversity, though it is worth noting that locus-specific estimates of *π*_b_ can be unreliable and should be treated with caution (Hahn, 2018). For clarity, we use ‘differentiation’ to refer to *F* _ST_ and ‘divergence’ to refer to *π*_b_ following Ravinet *et al*. (2017).

To visualize the distribution of putatively informative loci across the genome, Manhattan plots were created using the qqman package (Turner, 2018) in R for the mean decrease in Gini Index from random forest, the posterior inclusion probabilities from BSLMM, *F* _ST_, and *π*_b_. Additionally, the strength of linkage disequilibrium (LD) across the genome was quantified separately for autosomes and the Z chromosome using the the squared correlation coefficient (*r*^2^; Hill & Robertson, 1968) as calculated using the *--geno-r2* option in vcftools. The physical extent of LD was determined as the distance at which *r*^2^ fell below 0.2, and the shape of the LD curve was estimated using nonlinear regression using the *nls*function in R following the equations described by Remington *et al*. (2001), which were based on earlier work by Sved (1971) and Hill & Weir (1988).

## Results

Sequencing yielded a total of 619,374,339 reads, of which 412,722,330 were retained following preliminary filtering. Of the 435,753 variant sites, 8,366 were retained after 1) removing an individual that sequenced poorly; 2) filtering sites with minor allele frequency less than 0.02 and missing data for more than 70% of individuals; and 3) retaining only one locus per scaffold. After removing an additional 144 loci with mean depth > 35 and 6 loci with *F* _IS_ < –0.5 as part of an effort to exclude paralogous sites, the final dataset consisted of 479 individuals and 8,222 loci with an average depth of 12.1X per locus per individual.

Across the five replicate entropy chains, effective sample sizes were greater than 400 and potential scale reduction factors were less than 1.02, suggesting good chain mixing and convergence. While ancestry classifications were broadly consistent with wing morphology for individuals that were identified as *C. eriphyle* or *C. eurytheme* (94.5% and 99.3% concordance, respectively), only 17 of the 157 phenotypic hybrids had admixture proportions between 0.1 and 0.9 (Table S2). The majority of genetic hybrids were sampled in 1999 (both F_1_ hybrids and backcrosses), though a few backcrosses were also sampled in 2000 (Fig. 2). Additionally, one individual was sampled in 2001 that had *q* = 0.5 and *Q*_12_ = 0, a combination that is consistent with later generation (F_N_) hybrids. As for the alba females, 14 were classified as *C. eriphyle*, 59 as *C. eurytheme*, and 2 as hybrids (Table S2). Patterns of ancestry were broadly consistent for the full analysis (Fig. 2) and the analysis considering only males (Fig. S1).

In addition to considering all phenotypically hybrid (Hybrid_ALL_) and alba (Alba_ALL_) individuals as their own categories in the random forest models, we also considered split categories of these individuals based on the admixture proportion results from entropy (Hybrid_ER_ *q <* 0.1; Hybrid_HY_ 0.1 *< q <* 0.9; Hybrid_EU_ 0.9 *< q*; Alba_ER_ *q <* 0.5; Alba_EU_ *q >* 0.5). Random forest was only able to accurately assign individuals for models that directly compared genetically *C. eurytheme* and *C. eriphyle* individuals, irrespective of the morphological phenotype. Out of bag (OOB) errors were 0% for comparisons of orange versus yellow, genetically *eurytheme* versus *eriphyle* albas (Alba_EU_; Alba_ER_), genetically *eurytheme* versus *eriphyle* hybrids (Hybrid_EU_; Hybrid_ER_), and additional combinations thereof (Table 1). Specifically for the orange versus yellow comparison (based on morphology), most of the putatively causal loci (those with the greatest mean decrease of the Gini index) resided on the Z chromosome, but many were also found on the autosomes (e.g., chromosomes 2, 8, 13, 19; Fig. 3). Models were unable to parse alba females from other individuals from the same genetic background (Alba_EU_ versus Orange OOB = 28.57%; Alba_ER_ versus Yellow OOB = 12.70%). Similarly, morphologically hybrid yet genetically non-hybrid individuals could not be distinguished from other individuals from the same genetic background (Hybrid_EU_ versus Orange OOB = 32.18%; Hybrid_ER_ versus Yellow OOB = 36.78%). Genetically hybrid individuals could not be differentiated from either orange and yellow individuals (OOB = 7.27%) or Hybrid_EU_ and Hybrid_ER_ individuals (OOB = 8.28%). Finally, there were no detectable differences across the five years when considering all individuals (OOB = 54.70%), a *eurytheme* subset (OOB = 70.07%), or a *eriphyle* subset (OOB = 50.91%).

**Figure 3:**
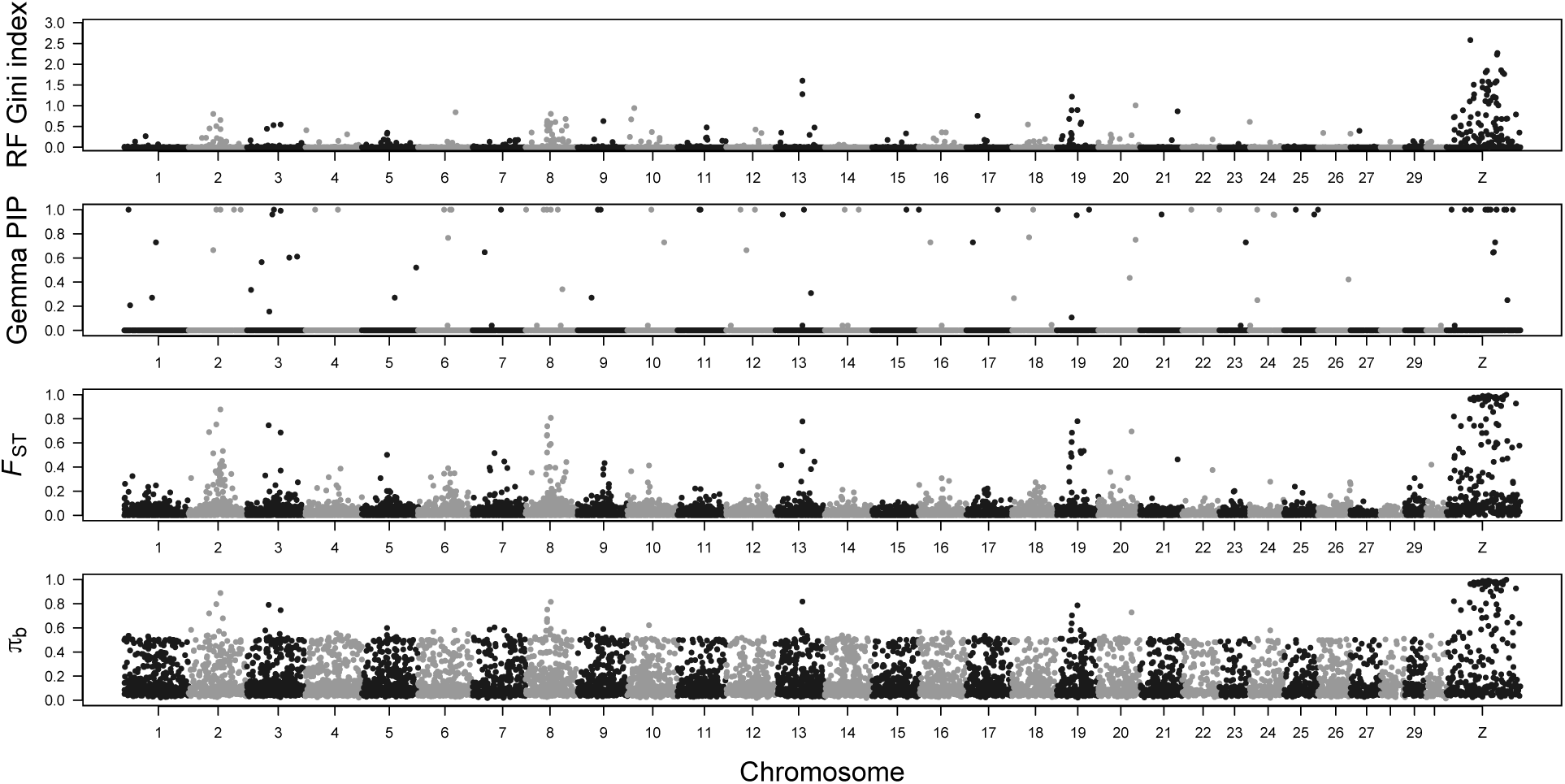
While many of the loci putatively associated with differences between *C. eriphyle* (yellow) and *C. eurytheme* (orange) individuals are concentrated on the Z chromosome, there are still many implicated sites found on several autosomes. Manhattan plots show locus-specific estimates of the mean decrease in the Gini index from random forest, posterior inclusion probability (PIP) from gemma’s posterior inclusion probability (PIP), *F* _ST_, and *π*_b_. These four complementary metrics quantify genomic differences between the two species. The Gini index (from random forest) and PIP (from gemma) measure the strength of association between genomic regions and the two phenotypes (orange versus yellow) in formal statistical prediction models, whereas *F* _ST_ and *π*_b_ are direct transformations of allele frequencies of individuals with the two phenotypes. Please note that these comparisons were performed for morphologically identified *C. eurytheme* and *C. eriphyle* individuals only; intermediate individuals were not included.

**Table 1:** Out of bag errors for the random forest models considered in this study. Response categories with classification errors < 0.05 are bolded. For color models, orange refers to *C. eurytheme* and yellow refers to *C. eriphyle*. Individuals identified as alba based on morphology were either lumped together (Alba_ALL_) or split based on the admixture proportion (*q*) from entropy (Alba_ER_: *q* < 0.5; Alba_EU_: *q* > 0.5). Similarly, individuals identified as hybrid based on morphology were either lumped together (Hybrid_ALL_) or split based on admixture proportion (Hybrid_ER_: *q* < 0.1; Hybrid_HY_: 0.1 < 0.9; Hybrid_EU_: 0.9 < *q*). Year models were run using either all, only orange, or only yellow individuals.

| Response | Response categories | Out of bag error |
| --- | --- | --- |
| Color | **Orange** + **Yellow** | 0.00% |
|  | **Orange** + **Alba<sub>ER</sub>** | 0.00% |
|  | **Orange** + **Hybrid<sub>ER</sub>** | 0.00% |
|  | **Yellow** + **Alba<sub>EU</sub>** | 0.00% |
|  | **Yellow** + **Hybrid<sub>EU</sub>** | 0.00% |
|  | **Alba<sub>EU</sub>** + **Alba<sub>ER</sub>** | 0.00% |
|  | **Hybrid<sub>EU</sub>** + **Hybrid<sub>ER</sub>** | 0.00% |
|  | **Orange** + **Yellow** + Hybrid<sub>HY</sub> | 7.27% |
|  | **Hybrid<sub>EU</sub>** + **Hybrid<sub>ER</sub>** + Hybrid<sub>HY</sub> | 8.28% |
|  | **Yellow** + Alba<sub>ER</sub> | 12.70% |
|  | **Orange** + **Yellow** + Alba<sub>ALL</sub> | 23.29% |
|  | **Orange** + Alba<sub>EU</sub> | 28.57% |
|  | **Orange** + Hybrid<sub>EU</sub> | 32.18% |
|  | **Orange** + Yellow + Hybrid<sub>ALL</sub> | 34.41% |
|  | **Yellow** + Hybrid<sub>ER</sub> | 36.78% |
|  | **Orange** + Yellow + Hybrid<sub>ALL</sub> + Alba<sub>ALL</sub> | 45.93% |
| Year | 1997<sub>ER</sub> + 1998<sub>ER</sub> + 1999<sub>ER</sub> + 2000<sub>ER</sub> + 2001<sub>ER</sub> | 50.91% |
|  | 1997<sub>ALL</sub> + 1998<sub>ALL</sub> + 1999<sub>ALL</sub> + 2000<sub>ALL</sub> + 2001<sub>ALL</sub> | 54.70% |
|  | 1997<sub>EU</sub> + 1998<sub>EU</sub> + 1999<sub>EU</sub> + 2000<sub>EU</sub> + 2001<sub>EU</sub> | 70.07% |

For the orange versus yellow wing color phenotype, the proportion of variance in phenotypes explained (PVE) and the proportion of genetic variance explained (PGE) by the BSLMM of gemma were both effectively 1, suggesting near-perfect heritability of wing color (narrow-sense heritability *h*^2^ = PVE x PGE; Bresadola *et al*., 2019; McFarlane & Pemberton, 2021). While several of the loci with high posterior inclusion probabilities (PIPs) were concentrated on the Z chromosome, a greater proportion of high PIP loci were found on the autosomes relative to the results from random forest (Fig. 3). It is worth noting that a simulation study (Jahner *et al*., 2024) found relatively high false positive rates for BSLMM relative to other comparable sparse modeling approaches, particularly when individual sample sizes are low, suggesting care is needed when evaluating candidates identified by this approach.

For individuals that were morphologically identified as orange and yellow, mean genome-wide differentiation and divergence was relatively low (*F* _ST_ = 0.040; *π*_b_ = 0.161) except for a subset of loci with extreme values. As in the other analyses, many but not all of the loci with elevated values resided on the Z chromosome (Fig. 3), where mean levels of differentiation and divergence were much higher than on the autosomes (Z chromosome *F* _ST_ = 0.278; *π*_b_ = 0.316; autosomes *F* _ST_ = 0.032; *π*_b_ = 0.156). Estimates of LD decay (based on all individuals) differed by several orders of magnitude between the Z chromosome (*r*^2^ declines below 0.2 at 274,650 bp) and the autosomes (max *r*^2^ = 0.061 at 100 bp). Importantly, the physical extent of LD decline on the Z chromosome was roughly seven times larger than the genome-wide marker density (1 locus per 39 kbp).

## Discussion

The 22 year record of parental abundance and hybridization at our focal location is characterized by highs and lows of counts that vary by orders of magnitude (Fig. 2; Jahner *et al*., 2012). The five years studied here, from 1997 to 2001, were chosen to capture one of the highest years for hybridization (based on wing phenotypes), as well as a rebound from an extreme low in density for one of the species. In response to drought conditions in 1994, farmers in the region of our study site allowed cultivated alfalfa to senesce in the latter part of summer. As a consequence, densities of *C. eurytheme* and *C. eriphyle* plummeted, and the latter was temporarily extirpated, with recolonization in 1996. Hybrid phenotypes were not observed during the period of extirpation, but frequencies of intermediate individuals rose again starting in 1997 and reached a peak in 1999 (Jahner *et al*., 2012). Consistent with that history, our genomic survey identified 1999 as the year with the most individuals with hybrid ancestry; zero hybrids were identified in the first two years, and very few in the latter two years (2000 and 2001; Fig. 2).

Individuals identified genetically as hybrids were almost always (a priori) classified as hybrids based on morphology (i.e., there were few false positives from the genetic perspective; Table S2). The opposite was not true: in many cases individuals identified as hybrids based on wing phenotype were assigned based on genomic composition to one parental species or the other. This contrast might be the result of error as the consequence of the subjective nature of morphological assignment as opposed to a quantitative assessment of color phenotypes (e.g., Hanly *et al*., 2023). On the other hand, it is also possible that morphological assignment of apparently-intermediate phenotypes was indeed accurate but reflects causal variants not captured by our sparse sampling of the genome (marker density = 1 locus every 39 kbp; Lowry *et al*., 2017). That likely possibility (important genetic variants not included in our dataset) is consistent with the fact that our sequencing did not capture genetic variants associated with alba (an unambiguous, genetically-based, white wing phenotype; Woronik & Wheat, 2017; Tunström *et al*., 2023) as revealed by relatively little success in random forest analyses attempting to predict alba individuals (Table 1). This could suggest a nuanced interpretation that apparently-intermediate phenotypes might have two sources: in some cases, they are true, first-generation hybrids; in other cases (perhaps the majority of cases), they are the result of color polymorphism segregating independently within the two parental species. The latter possibility might be the result of convergent evolution, past hybridization, or the maintenance of ancestral polymorphism, which was indeed the conclusion of Dwyer *et al*. (2015), albeit based on fewer loci without the power to detect true hybrids.

Regardless of the explanation for false positives in the morphological identification of hybrid individuals, the more unambiguous evolutionary conclusion from our results is a confirmation and quantification of hybridization in this system, and variability in introgression among years (Fig. 2). Pervasive hybridization, even if highly variable among years, is consistent with the mostly low differentiation across the genome for the two parental species (Fig. 3). The low differentiation between our focal taxa parallels the findings of Ficarrotta *et al*. (2022), comparing between *C. eurytheme* and *C. philodice*. In that case, differentiation was concentrated on the sex chromosome, with other instances of differentiation rarely rising to *F* _ST_ = 0.4. In contrast, we find many loci on autosomes with *F* _ST_ values above 0.5 and posterior inclusion probabilities at or near 1 in models for color variation (Fig. 3), though not on chromosome 18, which was previously implicated by Hanly *et al*. (2023). The differences between these studies, with a greater number of potential causally-linked autosomal variants in our study, could be related to the greater amount of wild variation encompassed; but that resolution will have to wait on future comparative work with greater geographic sampling.

Even with more autosomal loci potentially predictive of species differentiation in our study relative to previous studies, the *Colias* complex in North America is clearly dominated by differentiation on sex chromosomes. The importance of sex chromosomes has been observed in other butterflies (Sperling, 1994; Bastide *et al*., 2023; Xiong *et al*., 2023), and generally raises the question of how morphological variation is maintained on the landscape in the face of persistent hybridization. For the eastern pair of hybridizing species, *C. eurytheme* and *C. philodice*, there are apparently asymmetric fitness consequences associated with Z chromosomes from the different parental backgrounds interacting with autosomal genes (Grula & Taylor Jr, 1980). Whether or not similar or other mechanisms contribute to the maintenance of sex-linked variation between the western species is an area where further work is needed. We should also acknowledge the possibility that hybridization in these animals is far from equilibrium and that the role of stochastic drivers, including weather, is still underappreciated. Our use of replicate historical samples has been important because it is clear that any one of the five years studied in isolation would have given an incomplete picture of the system, either under– or overestimating the prevalence of hybridization. Including samples from many more years of the historical record (Fig. 2), especially with complete genome resequencing, would undoubtedly add new insights to our understanding of the dynamics in this system. However, it is possible that variability might be the rule rather than the exception (McFarlane *et al*., 2024), particularly for the factors that govern the coupling (or lack thereof) of phenotype and ancestry.

## Acknowledgments

We dedicate this paper to the memory of Ward B. Watt, who had the breadth of vision to study *Colias* from the community to the molecular level. We thank two anonymous reviewers for providing helpful suggestions for improving the text, and we thank Noah Adams for helping sort and organize individual butterflies. Analyses were performed on the University of Wyoming’s Advanced Research Computing Center and its Beartooth Computing Environment, Intel x86_64 cluster (https://doi.org/10.15786/M2FY47).

## Funding

This study was supported by California Agricultural Experiment Station Projects CA-D*-AZO-3994-H, “Climatic Range Limitation of Phytophagous Lepidopterans,” and CA-D-EVE-5203-H, “Dynamics of Hybridization in Sulphur Butterflies,” AMS, Principal Investigator. JPJ and SEM were supported by the Modelscape Consortium with funding from NSF (OIA-2019528). MLF was supported by NSF (DEB-2114793). Sequencing was supported by start up funds from the University of Nevada, Reno to TLP.

## Notes

### Competing Interest Statement

The authors have declared no competing interest.

### Summary of Updates

This version incorporates changes associated with the first round of peer review.

## References

1. Bastide H, López-Villavicencio M, Ogereau D, et al. (2023) Genome assembly of 3 Amazonian *Morpho* butterfly species reveals Z-chromosome rearrangements between closely related species living in sympatry. GigaScience, 12, giad033.

2. Bresadola L, Caseys C, Castiglione S, Buerkle CA, Wegmann D, Lexer C (2019) Admixture mapping in interspecific *Populus* hybrids identifies classes of genomic architectures for phytochemical, morphological and growth traits. New Phytologist, 223, 2076–2089.

3. Brien MN, Enciso-Romero J, Parnell AJ, et al. (2019) Phenotypic variation in *Heliconius erato* crosses shows that iridescent structural colour is sex-linked and controlled by multiple genes. Journal of the Royal Society Interface Focus, 9, 20180047.

4. Buerkle CA, Gompert Z (2013) Population genomics based on low coverage sequencing: how low should we go? Molecular Ecology, 22, 3028–3035.

5. Charlesworth B (1998) Measures of divergence between populations and the effect of forces that reduce variability. Molecular Biology and Evolution, 15, 538–543.

6. Cruickshank TE, Hahn MW (2014) Reanalysis suggests that genomic islands of speciation are due to reduced diversity, not reduced gene flow. Molecular Ecology, 23, 3133–3157.

7. Cuthill IC, Allen WL, Arbuckle K, et al. (2017) The biology of color. Science, 357, eaan0221.

8. Danecek P, Auton A, Abecasis G, et al. (2011) The variant call format and vcftools. Bioinformatics, 27, 2156–2158.

9. Dwyer HE, Jasieniuk M, Okada M, Shapiro AM (2015) Molecular evidence for hybridization in *Colias* (Lepidoptera: Pieridae): are *Colias* hybrids really hybrids? Ecology and Evolution, 5, 2865–2877.

10. Ebdon S, Mackintosh A, Hayward A, et al. (2021) The genome sequence of the clouded yellow, Colias crocea (Geoffroy, 1785). Wellcome Open Research, 6, 284.

11. Ellers J, Boggs CL (2003) The evolution of wing color: male mate choice opposes adaptive wing color divergence in *Colias* butterflies. Evolution, 57, 1100–1106.

12. Ficarrotta V, Hanly JJ, Loh LS, et al. (2022) A genetic switch for male UV iridescence in an incipient species pair of sulphur butterflies. Proceedings of the National Academy of Sciences USA, 119, e2109255118.

13. Fumagalli M, Vieira FG, Korneliussen TS, et al. (2013) Quantifying population genetic differentiation from next-generation sequencing data. Genetics, 195, 979–992.

14. Gompert Z, Lucas LK, Buerkle CA, Forister ML, Fordyce JA, Nice CC (2014) Admixture and the organization of genetic diversity in a butterfly species complex revealed through common and rare genetic variants. Molecular Ecology, 23, 4555–4573.

15. Grula JW, Taylor Jr OR (1980) Some characteristics of hybrids derived from the sulfur butterflies, Colias eurytheme and C. philodice: phenotypic effects of the X-chromosome. Evolution, 34, 673–687.

16. Hahn MW (2018) Molecular population genetics. Oxford University Press.

17. Hanly JJ, Francescutti CM, Loh LS, et al. (2023) Genetics of yellow-orange color variation in a pair of sympatric sulphur butterflies. Cell Reports, 42, 112820.

18. Hill W, Robertson A (1968) Linkage disequilibrium in finite populations. Theoretical and Applied Genetics, 38, 226–231.

19. Hill W, Weir B (1988) Variances and covariances of squared linkage disequilibria in finite populations. Theoretical Population Biology, 33, 54–78.

20. Hudson RR, Slatkin M, Maddison WP (1992) Estimation of levels of gene flow from DNA sequence data. Genetics, 132, 583–589.

21. Jahner JP, Buerkle CA, Gannon DG, et al. (2024) Interpretable and predictive models based on high-dimensional data in ecology and evolution. bioRxiv, doi:10.1101/2024.03.15.585297.

22. Jahner JP, Shapiro AM, Forister ML (2012) Drivers of hybridization in a 66-generation record of *Colias* butterflies. Evolution, 66, 818–830.

23. Langmead B, Salzberg SL (2012) Fast gapped-read alignment with Bowtie 2. Nature Methods, 9, 357.

24. Li H, Durbin R (2009) Fast and accurate short read alignment with Burrows–Wheeler transform. Bioinformatics, 25, 1754–1760.

25. Li H, Handsaker B, Wysoker A, et al. (2009) The sequence alignment/map format and SAMtools. Bioinformatics, 25, 2078–2079.

26. Liaw A, Wiener M (2002) Classification and regression by randomForest. R News, 2, 18–22.

27. Lindtke D, Gompert Z, Lexer C, Buerkle CA (2014) Unexpected ancestry of populus seedlings from a hybrid zone implies a large role for postzygotic selection in the maintenance of species. Molecular Ecology, 23, 4316–4330.

28. Lowry DB, Hoban S, Kelley JL, et al. (2017) Breaking RAD: An evaluation of the utility of restriction site-associated DNA sequencing for genome scans of adaptation. Molecular Ecology Resources, 17, 142–152.

29. Martin SH, Dasmahapatra KK, Nadeau NJ, et al. (2013) Genome-wide evidence for speciation with gene flow in *Heliconius* butterflies. Genome Research, 23, 1817–1828.

30. McFarlane SE, Jahner JP, Lindtke D, Buerkle CA, Mandeville EG (2024) Selection leads to remarkable variability in the outcomes of hybridisation across replicate hybrid zones. Molecular Ecology, 33, e17359.

31. McFarlane SE, Pemberton JM (2021) Admixture mapping reveals loci for carcass mass in red deer x sika hybrids in Kintyre, Scotland. G3, 11, jkab274.

32. Monteiro A, Prudic KM (2010) Multiple approaches to study color pattern evolution in butterflies. Trends in Evolutionary Biology, 2, e2–e2.

33. Nielsen R, Paul JS, Albrechtsen A, Song YS (2011) Genotype and SNP calling from next-generation sequencing data. Nature Reviews Genetics, 12, 443.

34. Parchman TL, Gompert Z, Mudge J, Schilkey FD, Benkman CW, Buerkle CA (2012) Genome-wide association genetics of an adaptive trait in lodgepole pine. Molecular Ecology, 21, 2991–3005.

35. R Core Team (2022) R: a language and environment for statistical computing. R Foundation for Statistical Computing, Vienna, Austria. URL https://www.R-project.org/.

36. Ravinet M, Faria R, Butlin R, et al. (2017) Interpreting the genomic landscape of speciation: a road map for finding barriers to gene flow. Journal of Evolutionary Biology, 30, 1450–1477.

37. Remington DL, Thornsberry JM, Matsuoka Y, et al. (2001) Structure of linkage disequilibrium and phenotypic associations in the maize genome. Proceedings of the National Academy of Sciences USA, 98, 11479–11484.

38. Rockman MV (2012) The QTN program and the alleles that matter for evolution: all that’s gold does not glitter. Evolution, 66, 1–17.

39. Rutowski R, Macedonia J, Morehouse N, Taylor-Taft L (2005) Pterin pigments amplify iridescent ultraviolet signal in males of the orange sulphur butterfly, Colias eurytheme. Proceedings of the Royal Society B: Biological Sciences, 272, 2329–2335.

40. Sahara K, Yoshido A, Traut W (2012) Sex chromosome evolution in moths and butterflies. Chromosome Research, 20, 83–94.

41. Shapiro AM (2012) The correspondence between John Gerould and William Hovanitz and the evolution of the *Colias* hybridization problem (Lepidoptera: Pieridae). The Journal of Research on the Lepidoptera, 45, 27–37.

42. Shastry V, Adams PE, Lindtke D, et al. (2021) Model-based genotype and ancestry estimation for potential hybrids with mixed-ploidy. Molecular Ecology Resources, 21, 1434–1451.

43. Sperling FA (1994) Sex-linked genes and species differences in Lepidoptera. The Canadian Entomologist, 126, 807–818.

44. Sved J (1971) Linkage disequilibrium and homozygosity of chromosome segments in finite populations. Theoretical Population Biology, 2, 125–141.

45. Tunström K, Woronik A, Hanly JJ, et al. (2023) Evidence for a single, ancient origin of a genus-wide alternative life history strategy. Science Advances, 9, eabq3713.

46. Turner S (2018) qqman: an R package for visualizing gwas results using QQ and Manhattan plots. Journal of Open Source Software, 3, 731.

47. Wheat CW, Watt WB (2008) A mitochondrial-DNA-based phylogeny for some evolutionary-genetic model species of *Colias* butterflies (Lepidoptera, Pieridae). Molecular Phylogenetics and Evolution, 47, 893–902.

48. Woronik A, Wheat CW (2017) Advances in finding Alba: the locus affecting life history and color polymorphism in a *Colias* butterfly. Journal of Evolutionary Biology, 30, 26–39.

49. Xiong T, Tarikere S, Rosser N, Li X, Yago M, Mallet J (2023) A polygenic explanation for Haldane’s rule in butterflies. Proceedings of the National Academy of Sciences, 120, e2300959120.

50. Zhang J, Cong Q, Shen J, Opler PA, Grishin NV (2019) Genomics of a complete butterfly continent. bioRxiv, doi:10.1101/829887.

51. Zhou X, Carbonetto P, Stephens M (2013) Polygenic modeling with Bayesian sparse linear mixed models. PLoS Genetics, 9, e1003264.

