## Supplementary material for "Temporal dynamics of color polymorphism and hybridization in *Colias* butterflies": supp

| Phenotype | 1997 | 1998 | 1999 | 2000 | 2001 | Total |
|---|---|---|---|---|---|---|
| *C. eriphyle* | 8 | 21 | 28 | 25 | 29 | 111 |
| *C. eurytheme* | 30 | 27 | 26 | 29 | 25 | 137 |
| Hybrid | 11 | 18 | 71 | 30 | 27 | 157 |
| Alba | 15 | 15 | 15 | 15 | 15 | 75 |

Table S1: The number of individuals sequenced per morphological phenotype and collection year.

| | Ancestry classification | | |
|---|---|---|---|
| Phenotype | *C. eriphyle* | Hybrid | *C. eurytheme* |
| *C. eriphyle* | 104 | 6 | 0 |
| Hybrid | 74 | 17 | 66 |
| *C. eurytheme* | 0 | 1 | 136 |
| Alba | 14 | 2 | 59 |

Table S2: A confusion matrix of individual phenotypes based on wing morphology and genotypes based on admixture proportions (*C. eriphyle*: $q < 0.1$; Hybrid: $0.1 < q < 0.9$; *C. eurytheme*: $q > 0.9$).

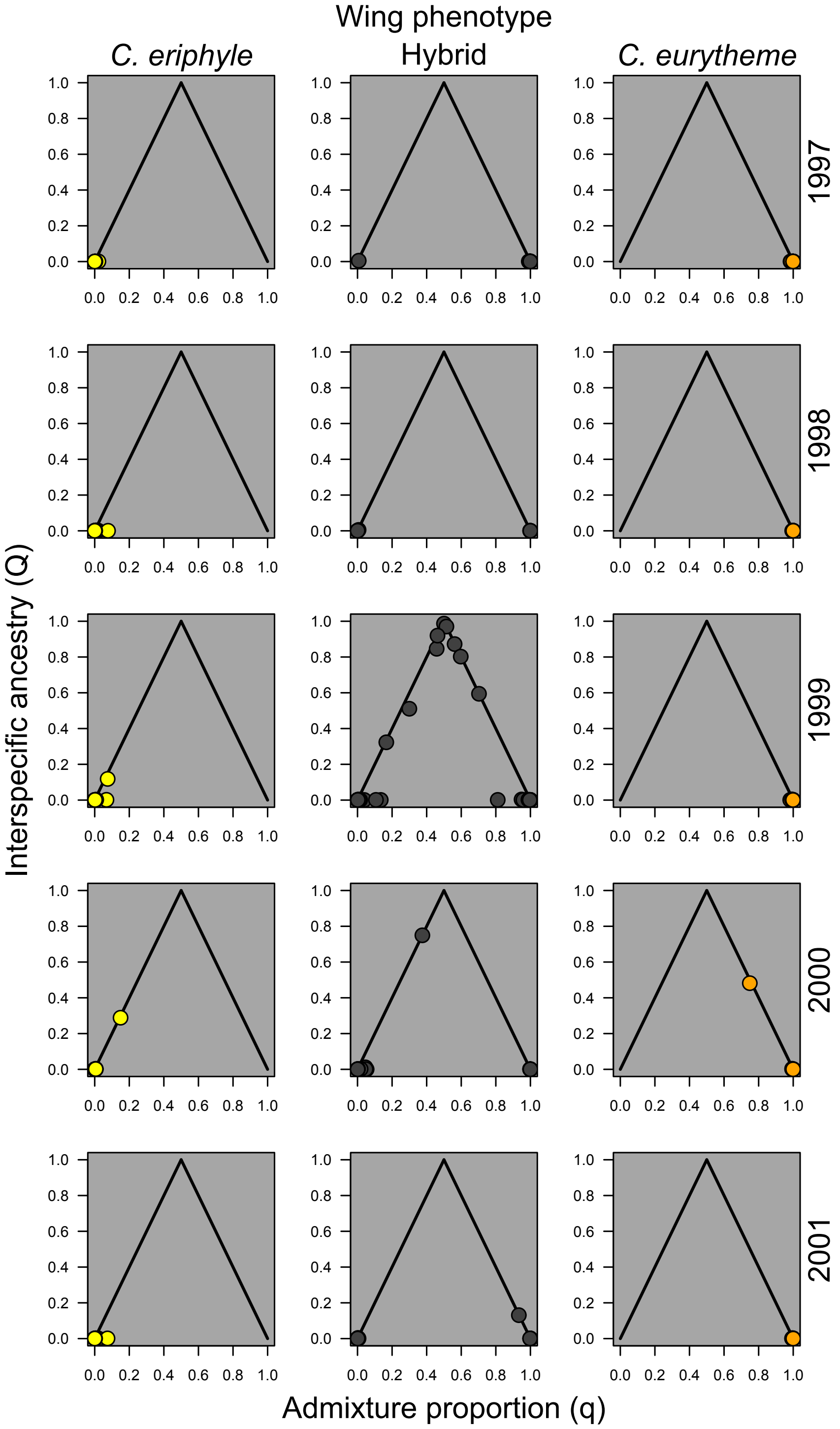

Figure S1: For all wing phenotypes and sampling years, individual ancestry was assessed for only males based on admixture proportion ($q$) and interspecific ancestry ($Q_{12}$) from `entropy`. Individuals with parental ancestry have $Q_{12} = 0$ and either $q = 0$ (*C. eriphyle*) or $q = 1$ (*C. eurytheme*); $F_1$ hybrids have $q = 0.5$ and $Q_{12} = 1$; $F_2$ hybrids have $q = 0.5$ and $Q_{12} = 0.5$ on average; and backcrosses have combinations of $q$ and $Q_{12}$ that reside on the solid black lines. See Fig. 2 in the main text for ancestry results from all individuals.
